# Inferring transmission routes for foot-and-mouth disease virus within a cattle herd using approximate Bayesian computation

**DOI:** 10.1101/2022.10.14.512099

**Authors:** John Ellis, Emma Brown, Claire Colenutt, David Schley, Simon Gubbins

## Abstract

To control an outbreak of an infectious disease it is essential to understand the different routes of transmission and how they contribute to the overall spread of the pathogen. With this information, policy makers can choose the most efficient methods of detection and control during an outbreak. Here we demonstrate a method for assessing the contribution of different routes of transmission using approximate Bayesian computation with sequential Monte Carlo sampling (ABC-SMC). We apply this to infer parameters of an individual based model of within-herd transmission of foot-and-mouth disease virus (FMDV), incorporating transmission through direct contact and via environmental contamination. Additionally, we use ABC-SMC for model selection to assess the plausibility of either transmission route alone being responsible for all infections. We show that direct transmission likely contributes the majority of infections during an outbreak of FMD but there is an accumulation of environmental contamination that can cause infections within a farm and also have the potential to spread between farms via fomites.

## 1 Introduction

Direct contact between infected and susceptible members of a population is the primary route of transmission for many viruses [26, 31, 39]. However, viruses can often persist on surfaces allowing the opportunity for transmission via fomites [4, 6, 13, 41]. Even if the contribution of environmental transmission is small during a local outbreak, persistence in the environment can contribute to long distance transmission via fomites and cause a resurgence if picked up by a new susceptible individual.

Determining the role of different routes of transmission of a virus is essential to have an effective control strategy during an outbreak. Therefore, there are many examples of virus studies that assess the routes of transmission, such as influenza [5, 33], coronaviruses such as MERS-CoV, SARS-CoV-1 and SARS-CoV-2 [29, 36, 53, 54], and animal viruses such as lumpy skin disease virus [8, 37], bluetongue virus [45], and foot-and-mouth disease virus (FMDV) [6, 13, 14, 20]. Some studies have used mathematical models to estimate the contribution of different routes of transmission. This requires parameterisation of the model, which can be achieved by using estimates from the literature [2, 35, 44], or by fitting different sets of parameters or different models to data [37, 53, 54, 45].

Foot-and-mouth disease (FMD) is a highly infectious disease that is known to be transmitted in a number of ways. FMDV infects cloven-hoofed animals such as cattle, sheep, goats, pigs and various wildlife species [22] and is shed from infected animals through their excretions and secretions, potentially remaining infectious for a prolonged period of time (depending on environmental conditions) [6, 13, 14]. This allows the virus to be carried over long distances via the wind or fomites, driving between farm transmission [6, 13, 14, 20]. Due to the fast rates of infection within a farm, the focus from authorities is to stop between-farm transmission and therefore previous models of FMDV epidemics usually treat each farm as an individual unit [17, 18, 32, 48]. This is a useful approach when responding to an outbreak but it simplifies the within-herd dynamics which can show how the infectiousness of individual farms changes over time and be used to assess surveillance strategies. Some studies model individual animals to calculate the infectiousness profile of an infected farm over time and estimate the detection time [9, 11, 47] but do not consider the accumulation of virus in the environment.

Here, we present the methodology and results from an individual based model of FMDV transmission within a cattle herd. Within the model, infected animals can pass on the virus directly and also shed virus into the environment, causing indirect transmission. The within-host infection dynamics are modelled to determine the rate of viral accumulation in the environment and the probability of direct transmission within the herd. We parameterise the model using approximate Bayesian computation sequential Monte Carlo (ABC-SMC) [23, 40, 50], fitting to data from the 2007 UK epidemic [42]. Using ABC-SMC with model selection, we then compare different scenarios where either one or both routes of transmission cause infection. By obtaining the number of animals becoming infected through different routes and finding the transmission model with best fit, we assess the importance of the different routes of transmission in an outbreak. Other model outputs include the level of environmental contamination and total infectiousness of animals in the herd over time. We also consider how the different routes of transmission influence summary measures such as the herd generation time and the proportion of infectiousness that occurs from a farm before animals show clinical signs (useful for assessing the effectiveness of reactive control measures) [10, 19].

## 2 Methods

### 2.1 Mathematical model

We simulate the within-herd transmission of FMDV using an individual based model (IBM) where transmission can occur via direct contact and via environmental contamination. An IBM allows us to model variability of within-host dynamics amongst animals, such as the level of viral shedding at a given time since infection. Unlike in compartmental modelling, which assumes a constant force of infection for a given number of infected animals, we model dynamic infectiousness that peaks a certain number of days post-infection. Similarly, the contribution to environmental contamination from an individual animal at any given time is dependent on its level of viral shedding.

Our model is informed by data from transmission experiments to quantify transmission of FMDV through direct contact and environmental contamination [10, 13]. In these experiments, cattle were either exposed to an infected bovine or a room where an infected bovine had recently been housed. By taking swabs from the exposed cattle and the environment, the within-host dynamics, infection rates and the rates of contamination could be estimated.

After the introduction of an infectious animal into a herd or viral contamination into the environment, the probability of any susceptible animal becoming infected over a given time interval is calculated. The number of new infections within a time interval is then determined stochastically for each individual animal.

#### 2.1.1 Viral shedding

Upon infection, the infectiousness of an animal is proportional to the level of viral shedding. The viral titre in an infected animal typically increases exponentially after infection before reaching a peak after which it decays exponentially back to zero [13, 27]. We model this as

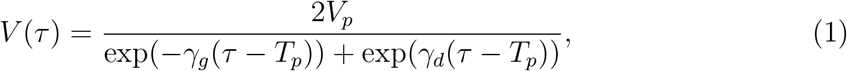

where *V_p_* is the level of peak titre, *T_p_* is the time of peak titre, *γ_g_* and *γ_d_* are the rates for the exponential viral growth and decay phases respectively and *τ* is the time since infection. The time taken for clinical signs to appear in an individual is also considered in the model, given by *T_c_*, and is linked to the shedding by assuming the times of peak titre and clinical onset are jointly distributed.

Parameters that determine the level of viral shedding are randomly generated to allow variation between individuals. *γ_g_*, *γ_d_* are generated from a gamma distribution with means 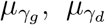 and shape parameters 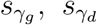 respectively. *V_p_* is generated from a log gamma distribution with parameters *μ_V_* and *s_V_* (the mean and shape parameter of the corresponding gamma distribution). The viral dynamics are linked to the onset of clinical disease by assuming the time of peak titre *T_p_* and incubation period *T_c_* follow a bivariate log normal distribution with parameters 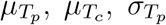 and 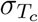 (the means and standard deviation on the log scale of the corresponding normal distribution) and a correlation coefficient *ρ_pc_*.

Based on transmission experiments, we assume that the level of viral shedding is proportional to the log viral titre, i.e.

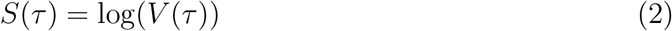

where *V*(*τ*) is given by Eqn. (1) and *S*(*τ*) is restricted to be non-negative.

#### 2.1.2 Environmental contamination

We model the level of environmental contamination such that it depends on the amount of virus shed by animals and the natural decay rate of virus in the environment [13]. The contamination and decay rates are assumed to vary between four areas: the floor, walls, trough and faeces. The choice of contamination areas arises from those sampled in the transmission experiment [13]. We use these terms broadly so that walls includes fence posts and floor includes straw bedding, for example.

The level of virus in each location is given by

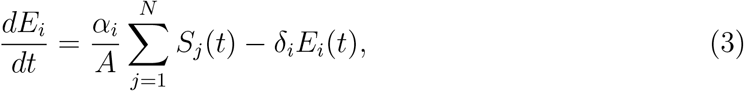

where *E_i_*, *i* = 1, …, 4 is the level of contamination found in the floor, walls, trough and faeces respectively. *α_i_* is the rate of contamination, *δ_i_* is the rate of decay, *N* is the population size of the herd and *A* denotes the relative size of the environment.

We assume homogeneous contamination in each area. This is supported by evidence from a contact study that there is a high level of spread of contaminants [34]. For example, faecal matter from a single cow-pat was detected on 80 ± 4% of cattle in a given group 12 hours later.

The rate of contamination is scaled by 1/*A* to account for the difference in size of the rooms used in the transmission experiments and the size of cattle housing on a farm. In a small environment, the contamination of virus in the environment is likely to be at a higher concentration than when spread over a larger farm environment. Intuitively, we would expect the size of an environment to be related to the size of the herd. Consequently, we assume that the environment size of all four areas is proportional to herd population and set *A* = *N* henceforth, unless otherwise stated. However, we note that the contamination rates are to be fitted to outbreak data, so if our estimate is unsuitable then the contamination rate will change to compensate.

#### 2.1.3 Transmission

Transmission of FMDV within the herd can occur through direct contact between animals or through environmental contamination, dependent on the level of viral shedding from infected animals and the level of virus in the environment. For direct transmission, the probability of an animal becoming infected via direct contact with an infected individual during the time interval [*t, t* + Δ*t*] is given by

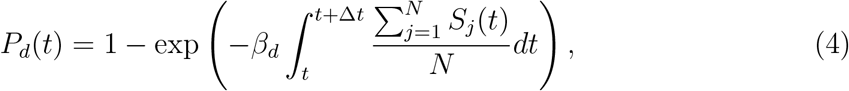

where *β_d_* is the direct transmission rate. As the sum of viral shedding is divided by the population size, this assumes homogeneous mixing between animals (contact between individuals is random) and frequency dependent transmission (the number of contacts is not dependent on herd size), which is appropriate for livestock diseases [31].

Similarly, the probability of a susceptible animal becoming infected due to environmental contamination in the interval [*t, t* + Δ*t*] is given by

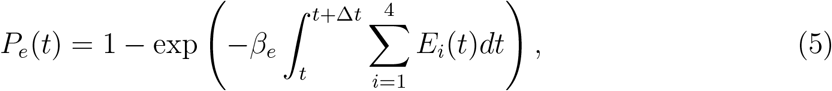

where *β_e_* is the environmental transmission rate which is assumed to be the same for all contaminated areas [13].

Therefore, the probability of a susceptible animal becoming infected in each time interval is given by

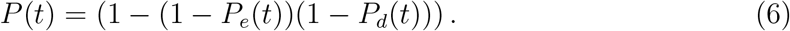

When the level of viral shedding, given by Eqn. (2), falls to zero, the animal is no longer infectious, i.e. its contribution to the rate of environmental contamination and the probability of direct transmission is zero. However, we do not consider recovery from clinical signs as we are only concerned with the time they first appear in each animal, given by *T_c_*. Lesions start to heal approximately 10 days after infection, though it can take several weeks for a full recovery by which time viral shedding has typically stopped [38, 43].

### 2.2 Data

Transmission experiments in high-containment animal facilities have been used to estimate transmission rates for FMDV. Susceptible cattle were either directly exposed to infected donor cattle [10] or placed in a room that had recently been occupied by infected cattle [13]. Nasal swabs were collected daily from donor cattle [10], swabs were taken from the environment[13], and the outcome of direct or environmental challenges recorded [10, 13]. These data were used to generate prior distributions for the shedding parameters, environmental contamination and decay rates and direct and environmental transmission rates.

To relate the results from transmission experiments to the field, we use data from the 2007 FMD epidemic in Surrey, UK, given in [42] (Table S1). During this epidemic, eight farms in Surrey and Berkshire in southern England were infected, resulting in the culling of 1578 animals, of which 278 were confirmed to be infected with FMDV. All of the cattle herds affected were small, with the largest herd containing 58 cattle. The lesion ages on five of the infected premises (IPs) were estimated prior to culling and are recorded in [42]. We refer to each IP using the nomenclature assigned by veterinary authorities and used in [42].

### 2.3 Model fitting

Model parameters (listed in Table S2) were estimated using approximate Bayesian computation sequential Monte Carlo (ABC-SMC) methods. ABC-SMC is a useful tool for stochastic models when it is difficult to define the likelihood and has been used previously with models of infectious diseases [3, 23, 24, 40, 50]. A goodness-of-fit measure that allows the simulation to be compared to the data is used to assess the model accuracy. The algorithm generates particles, or parameter sets, from a joint prior distribution and runs a model simulation for each. Particles are accepted when the simulation is within a certain distance from the data. Accepted particles then form an intermediate distribution with weights based on the prior probabilities. New particles are subsequently sampled based on their weights and perturbed in a new round with a new distance threshold. This continues until the goodness-of-fit measure is within an acceptable distance and the particles have converges to the posterior distribution.

The ABC-SMC algorithm was used to estimate parameters separately for each of the five IPs. For each IP, an outbreak is simulated in a herd with a population matching that given by the data. Informative prior distributions were obtained for all parameters using the data from transmission experiments (Table S2). A non-informative, uniform prior distribution was used for the number of animals infected at the start of the simulation. The model was fitted by comparing the daily appearance of clinical signs in model simulations with those estimated from lesion ages in Table S1. The goodness-of-fit measure was given by the residual sum of squares between simulated and observed daily incidence, i.e.

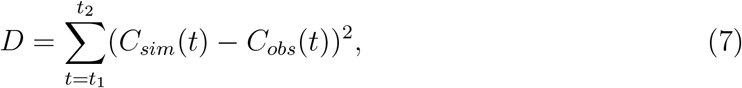

where *C_sim_*(*t*) is the simulated number of animals newly developing clinical signs on day *t*, *C_obs_*(*t*) is the number of cattle with new clinical signs on day *t* based on observed lesion ages, *t*_1_ is the day clinical signs first appeared in the herd and *t*_2_ is the day lesion ages were assessed (and the herd was culled). For each round, the algorithm was run until 10,000 particles were accepted. Convergence to the posterior distribution was assessed by visual inspection after each round. A Gaussian perturbation kernel was used with standard deviation equal to one tenth of the range of prior distribution.

ABC-SMC can also be used for selecting between multiple models [49, 50]. In this case an extra step is introduced whereby a model is selected according to the particle weights. This is then perturbed so that there is a probability for another model to be selected before a particle is generated from that model. We used the model selection algorithm to determine how models with direct transmission only or environmental transmission only compare to the combined direct and environmental transmission model. Both single transmission models are similar to the combined transmission model and the same prior distributions are used. In the direct transmission model, Eqns. (3) and (5) are removed and in the environmental transmission model, Eqn. (4) is removed.

We choose the prior probability distribution of each model to be uniform so that there is an equal chance of the three models to be selected in the initial round of the algorithm. After a model is selected, the probability of it being perturbed to a different model was 0.2 with a probability of 0.8 of it remaining as the selected model. The same goodness-of-fit measure was used as in Eqn. (7) and the particle perturbation kernel, number of particles accepted and assessment of convergence was implemented in the same way as with the single model parameter fitting. The output of the algorithm is the marginal posterior distribution of the model parameters which can be used to calculate the Bayes factors and determine the model with the best fit to the data [30, 49, 50].

### 2.4 Summary transmission measures

The outputs of the model include the amount of viral shedding by each animal over time and the levels of environmental contamination. The sum of all viral shedding by individual cattle and the amount of environmental contamination can be used to approximate the total infectiousness of an IP. We treat the infectiousness of viral shedding and environmental contamination separately but note that if the relative contribution towards between farm transmission was known, a combined total infectiousness could be calculated. The summary measures described in this section for combined infectiousness would likely be somewhere between those for shedding and environmental contamination.

A critical measure when examining reactive control of an infectious disease is the proportion of transmission that occurs before clinical signs or symptoms appear [10, 19]. From examining the within-herd dynamics of FMDV, a similar measure can be found for the farm scale, indicating the opportunity for infection between farms to occur before detection or control.

The proportion of transmission via direct contact before detection is given by

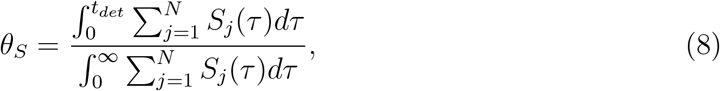

and the proportion of transmission via the environment before detection is given by

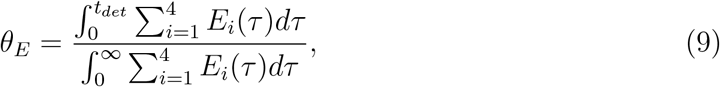

where *t_det_* is the time at which the herd is detected. If transmission is stopped at *t_det_*, then *θ_S_* and *θ_E_* can be used to show how much transmission is reduced by early detection.

Although it may be possible to combine transmission via direct contact and the environment to give a single figure for *θ*, it is not clear what the relative contribution of the two routes is likely to be and, hence, how this would be calculated. More knowledge of between-farm transmission dynamics is required to determine the significance of each transmission route. For example, environmental contamination may be more significant in between farm-transmission than within-farm transmission due to the transportation of fomites being a significant risk factor.

Total infectiousness can also be used to calculate the herd generation time, defined to be the interval between the infection time of an IP and the infection time of IPs arising from it [25]. Generation times can be estimated by contact-tracing and examining times of infection during an outbreak [25, 28], but it can also be calculated explicitly when the levels of infectiousness are known [21]. As our model contains two routes of transmission, herd generation times are obtained for direct and environmental transmission separately.

For direct transmission this is given by

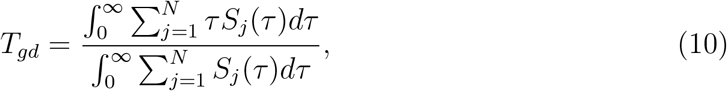

and for environmental transmission,

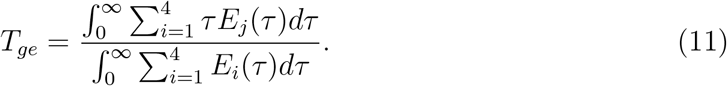

Comparing the two herd generation times can highlight the differences in the between-farm transmission dynamics that might arise from the two routes of transmission. This can also be useful in determining the cause of infection when the generation time is known. For example, if *T_g_* is closer to *T_ge_* than *T_gs_*, then it may be more likely that infection occurred via the transportation of fomites rather than the movement of animals.

## 3 Results

### 3.1 Parameter estimation

The posterior distributions for the parameters for each IP are shown in Figs. S1 and summarised in Table S2. In most cases the posterior distributions do not diverge greatly from the prior distributions. The 90% credible intervals for the posterior distributions are consistently larger than the prior, possibly due to the high degree of stochasticity in the model, meaning that a range of parameter sets can produce results closely matching the data. One notable difference is that for all IPs there is an increase in the direct transmission parameter (*β_d_*).

Posterior distributions are also largely similar between IPs with some variation in parameters relating to the time clinical signs develop and the direct transmission parameter. IP4b differs the most from other IPs in that it has a slightly higher mean peak titre (*μ_V_*), lower time for clinical signs to appear (controlled by 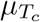 and 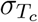) and a higher direct transmission parameter (*β_d_*) in particular. This is the largest herd, with 54 animals all of which were judged to have had lesions appear within 5 days of the first, suggesting the virus spread rapidly through the herd. At other IPs, not all animals had lesions at this point (Table S1).

The posterior distributions for the number of initially infected animals are similar for IP1b, IP2a, IP3b and IP4b (see Fig. 1), all of which had similar herd sizes (from 38 to 54), although it is slightly higher for IP3b and slightly lower for IP4b. The herd size on IP7 was smaller at 16 cattle and therefore there are fewer initial infections.

**Figure 1:**
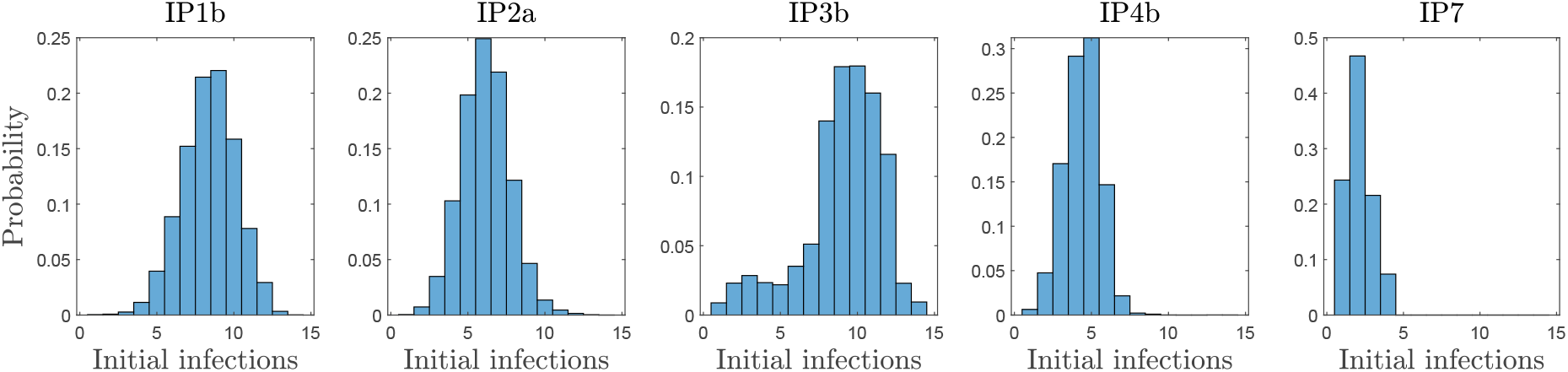
Posterior distributions for the number of animals initially infected with foot-and-mouth disease virus for five farms infected in the 2007 UK epidemic.

### 3.2 Transmission dynamics

A comparison of simulated and observed lesion ages and the cumulative number infections are shown in Fig. 2. Although the observed data occasionally fall outside of the 90% credible intervals, the simulated data follows the general pattern of the outbreaks, with the number of animals with clinical signs usually peaking three to four days after the first lesions appear. It should be noted that there is uncertainty in the accuracy of lesion ages which may explain differences between observed and simulated outbreaks, particularly for lesions more than five days old [42].

**Figure 2:**
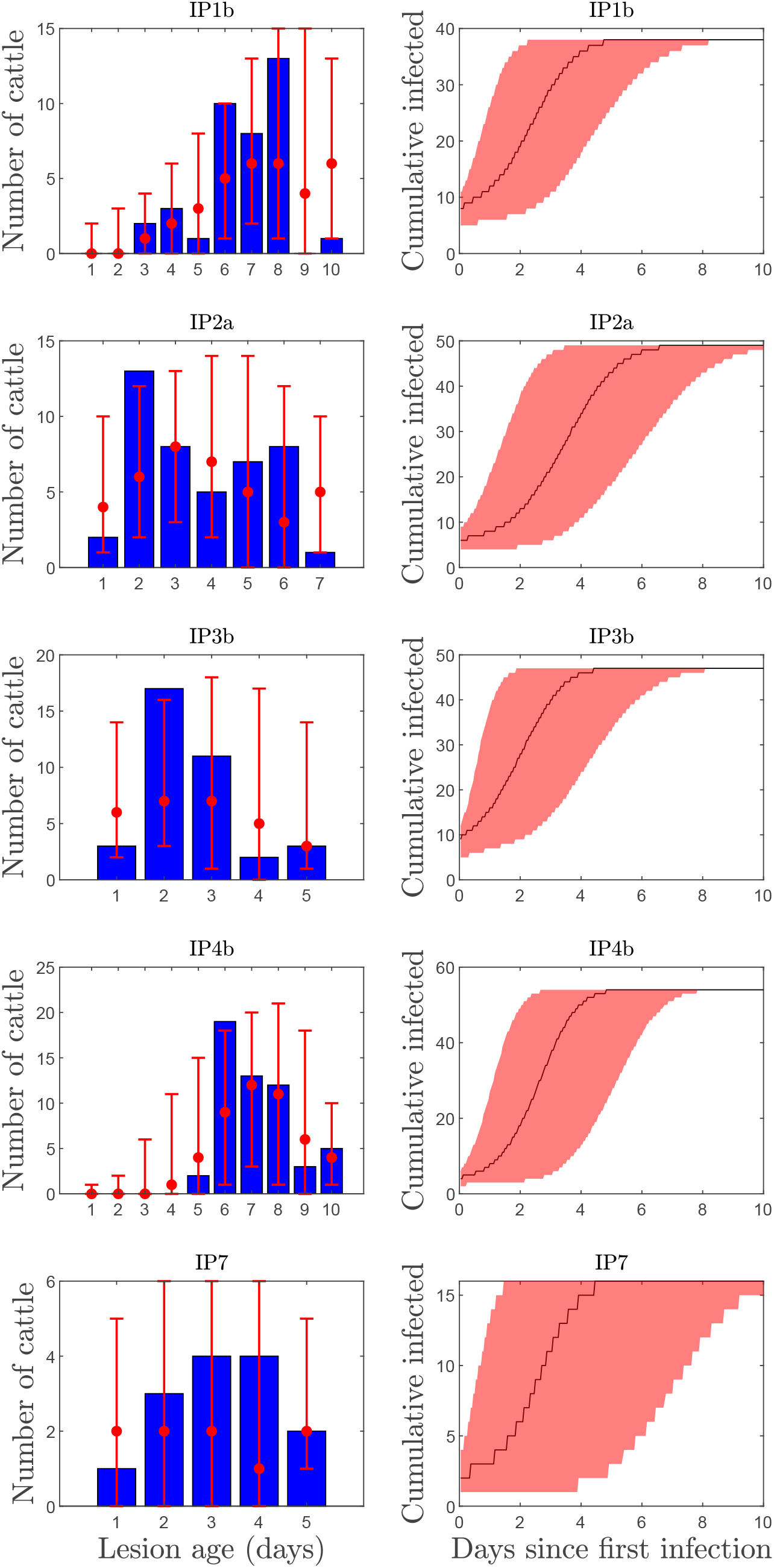
A comparison between observed and simulated cattle lesion ages (left) (red, showing the median and 90% credible interval) and lesion age estimations [42] (blue bars) and the median and 90% credible interval of the cumulative number of infected animals (right) for five farms infected with foot-and-1m1outh disease virus in the 2007 UK epidemic.

The cumulative number of infected cattle increases at similar rate for all IPs. The entire population usually becomes infected in two to eight days with a median of four to five days, although it takes slightly longer at IP2a and there is more variability for IP7.

### 3.3 Comparing routes of transmission

As shown in Fig. 3, the model predicts that transmission mostly occurs through direct contact and only infrequently through the environment. Infections through the environment also tend to occur later than those from direct contact. Infections via direct transmission peak two to three days into the outbreak, while those via the environment usually peak one day later (Fig. 4). Very few infections occur after day 10 although this is due to the small number of susceptible animals left at this time.

**Figure 3:**
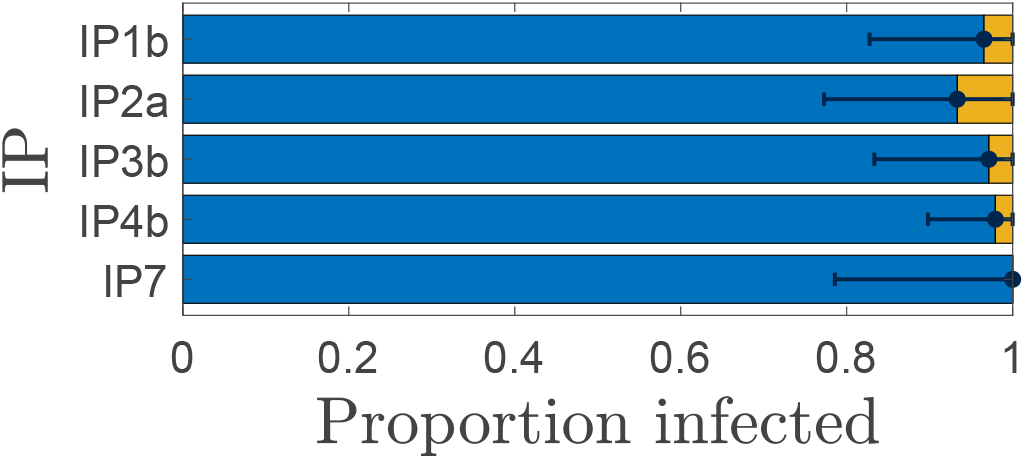
The proportion of animals infected with foot-and-mouth disease via direct contact (blue, left) or environmental transmission (yellow, right) on each of the five IPs. The marker where the two different coloured bars meet shows the median, and the error bars show the 90% credible interval.

**Figure 4:**
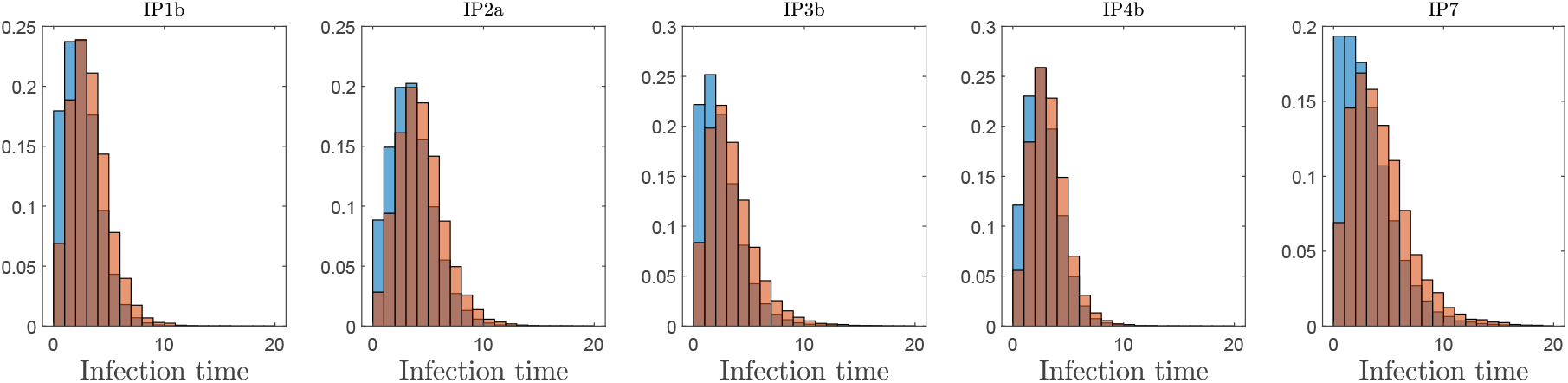
The distribution of times of infection via the direct transmission (blue) and environmental transmission (red) on each of the five IPs. The data has been normalised to show the proportion of infections from each route that happen within each day.

The output from ABC-SMC model selection, comparing models with either direct transmission, environmental transmission or both, shows that either the combined model or the direct transmission only models are the best choice to fit the data from all five IPs. The Bayes factors [30] for each model on each IP are shown in Table 1. The larger the Bayes factor, the stronger the evidence in favour of the model labelled first, e.g. *B_cd_* shows the strength of evidence in favour of the combined transmission model and against the direct transmission only model. The Bayes factor for combined and direct transmission (*B_cd_*) is close to one for all five IPs, indicating that there is no strong evidence in favour of either model. However when either is compared to the environmental transmission model, there is strong evidence that the combined or direct transmission models are a better fit. This is unsurprising as in the combined model the majority of infections arise via direct contact (see Fig. 3). In this case, the environmental only model can be discounted.

**Table 1:** Bayes factors for model selection. The models are combined transmission (*c*), direct transmission only (*d*) and environmental transmission only (*e*).

| | $B_{cd}$ | $B_{ce}$ | $B_{de}$ |
| --- | --- | --- | --- |
| IP1b | 1.02 | $1.29 \times 10^6$ | $1.22 \times 10^6$ |
| IP2a | 0.844 | $9.75 \times 10^4$ | $1.16 \times 10^5$ |
| IP3b | 1.70 | $9.18 \times 10^8$ | $5.39 \times 10^8$ |
| IP4b | 1.26 | $1.25 \times 10^{15}$ | $9.94 \times 10^{14}$ |
| IP7 | 0.871 | $2.78 \times 10^4$ | $3.19 \times 10^4$ |

### 3.4 Summary transmission measures

The total infectiousness of an IP from infected animals and environmental contamination can be approximated by the sum of viral shedding from all infected animals and the total amount of virus in the environment respectively, shown in Fig. 5. Although the two types of infectiousness cannot be compared directly as they contribute differently to transmission (also note the different scales on the y-axes), they show the difference in the shape of the curve, including the time when infectiousness peaks.

**Figure 5:**
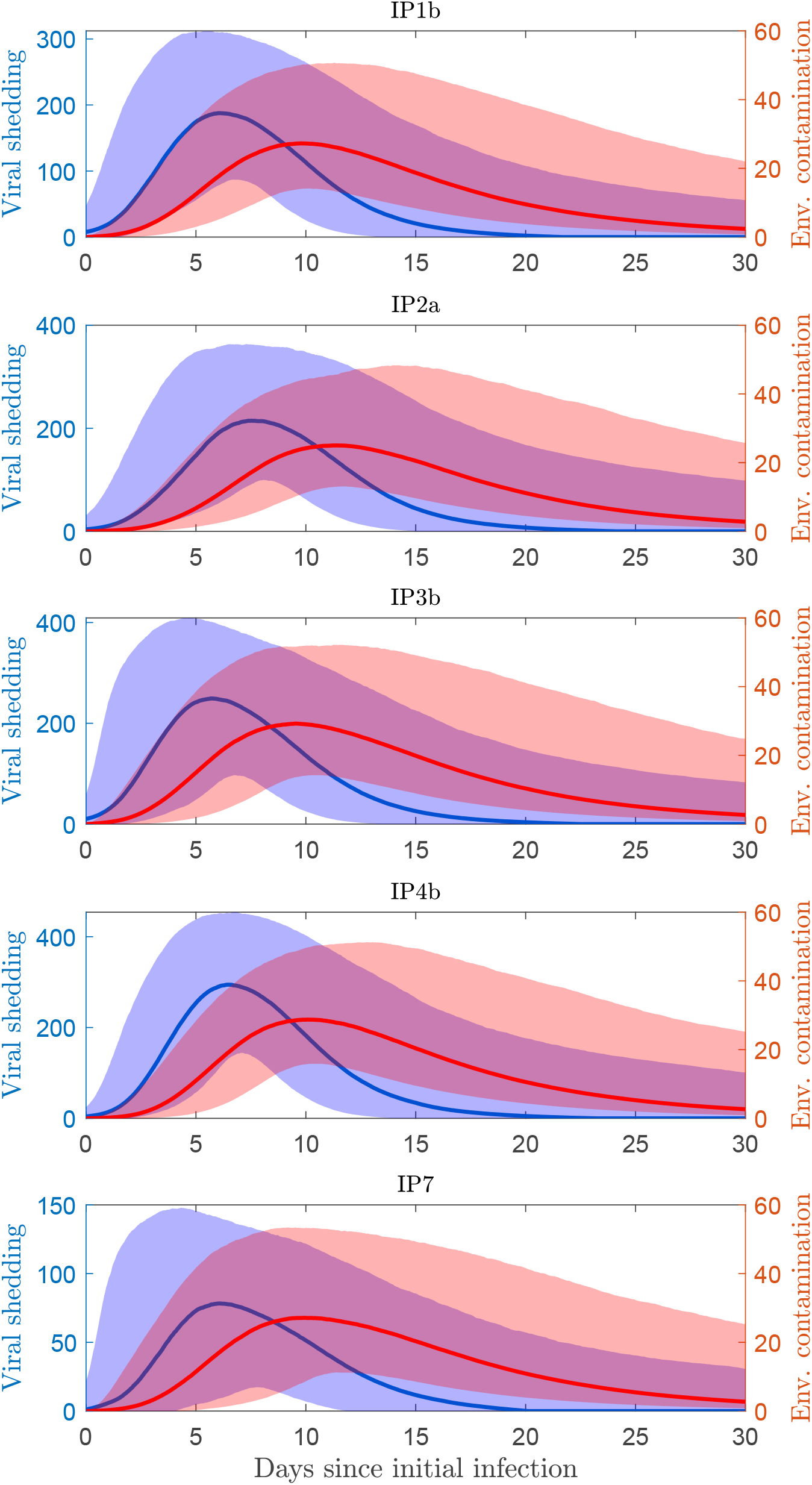
The total infectiousness of a farm infected with foot-and-mouth disease virus measured by the total shedding from infected cattle (blue, left axis) and the level of virus in the environment (red, right axis). The median is represented by a solid line and the shaded area is the 90% credible interval.

Total infectiousness can be used to assess the risk of between farm transmission. In particular, the proportion of transmission that occurs before detection (*θ_S_* and *θ_E_*) can demonstrate the importance of early detection and intervention. We assumed that a herd would be detected either a given number of days after clinical signs first appear or the day after a given proportion of animals show clinical signs. Results are shown in Fig 6. The proportion of transmission via the environment before detection is far less than the proportion of transmission via direct contact before detection for each time of detection. Even 10 days after the first clinical signs, approximately only half of environmental transmission has occurred. The proportion of transmission via direct contact before detection remains low (< 10%) if detected up to 1-2 days after the first clinical signs appear. If detected five days after the appearance of clinical signs, this proportion increases to over half in most cases and, if detected 10 days after, it has increased to approximately 90%.

**Figure 6:**
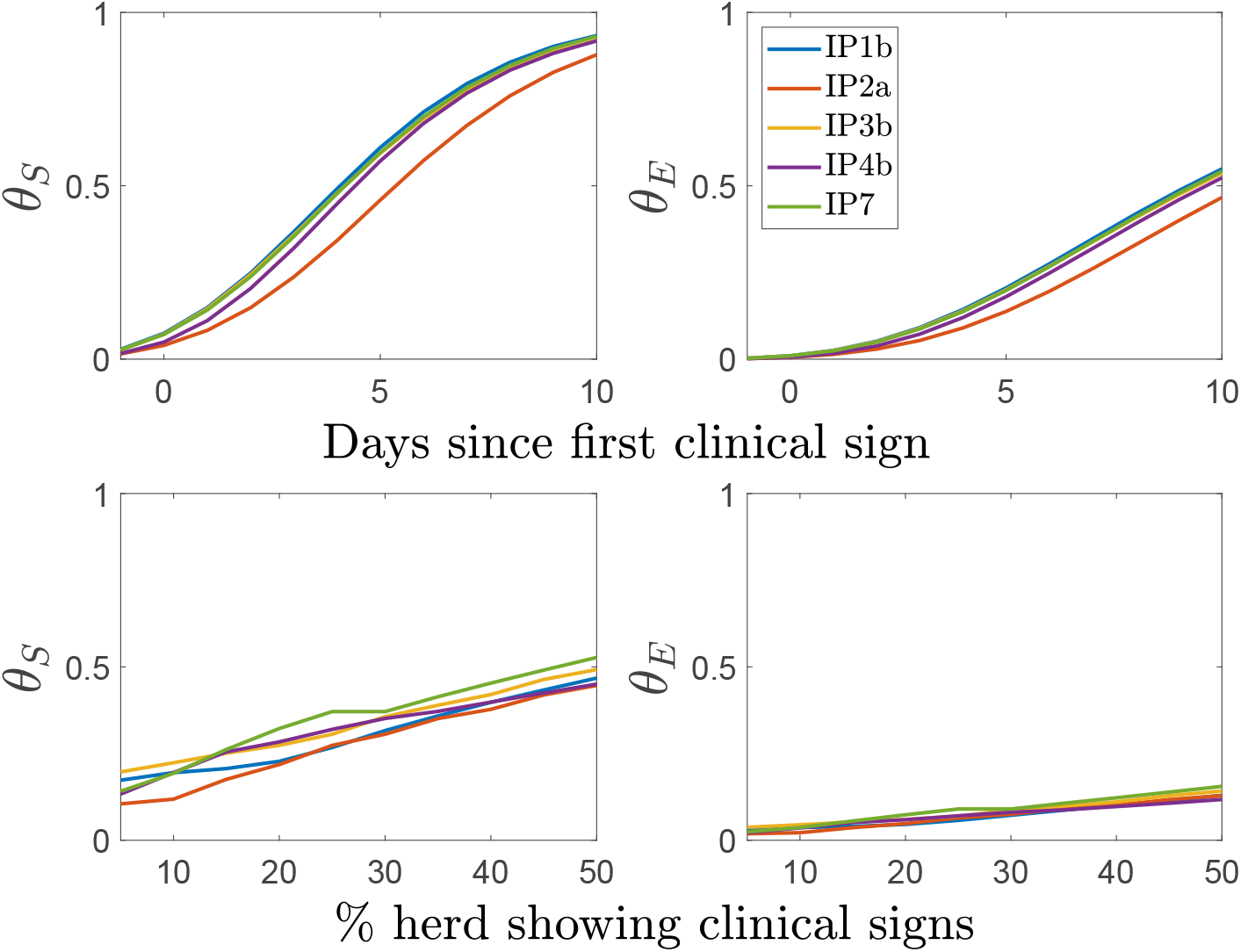
Proportion of infectiousness that has occurred before detection, assuming different detection times. Top row shows the number of days since the first animal to show clinical signs and the bottom row show *θ* one day after a given proportion of the herd have clinical signs.

Taking the times one day after a proportion of the herd shows clinical signs, we see that there is not a large difference in proportion of transmission before detection between the thresholds of 5% and 10% of animals. In some cases, there is not a large increase from the proportion of transmission that occurs one day after the first clinical signs to one day after 10% show clinical signs. There are more substantial rises from the 10% to 20% and 20% to 50% thresholds.

The sum of viral shedding and environmental contamination are also used to calculate the herd generation times, *T_g_*. For each IP, the herd generation times for direct and environmental transmission are given in Table 2. The median generation times for the IPs are all approximately 7-9 days for direct transmission and 13-15 days for environmental transmission.

**Table 2:** Median (90% credible interval) herd generation times (days) for viral shedding (*T_gd_*) and environmental contamination (*T_ge_*) for each IP.

|  | IP1b | IP2a | IP3b | IP4b | IP7 |
| --- | --- | --- | --- | --- | --- |
| $T_{gd}$ | 7.75<br>(4.97-15.9) | 9.07<br>(5.87-17.3) | 7.54<br>(4.72-16.1) | 8.12<br>(5.39-16.4) | 8.15<br>(4.48-17.5) |
| $T_{ge}$ | 13.8<br>(11.0-20.2) | 15.1<br>(11.9-22.4) | 13.7<br>(10.8-20.5) | 14.2<br>(11.4-20.5) | 14.1<br>(10.5-21.7) |

### 3.5 Sensitivity Analysis

#### 3.5.1 Environment size

In Eqn. (3), we introduced the size of the environment, *A*, which scales the rate of viral contamination in the environment. It is reasonable to assume that the level of contamination at a location will be dependent on the size of the whole environment, as cattle move around a premises. In the absence of more information, we assumed a linear relationship between the herd size and the size of the environment. In reality, there will be variation in the size of the environment where cattle are kept in depending on, for example, the farm, the type of cattle, herd management practices and the time of year. Here, we present the results when the size of the environment is not considered (the equivalent of setting *A* = 1 in Eqn. (3)) for IP1b (results for the other IPs are similar). The prior distributions are the same as in Table S2.

The posterior distributions for many of the parameters do not change when the environment is not scaled by herd size (Fig. S2). However, the mean peak titre (*μ_V_*) and both transmission parameters (*β_d_* and *β_e_*) have a slightly lower median than in the environmentally-scaled model which counteracts the impact of higher contamination driving more environmental transmission.

The level of environmental contamination is larger by a factor of more than 20 when *A* = 1 compared with when *A* = *N* (Fig. 7; cf. Fig. 2), and environmental transmission now contributes over half of all infections in most runs of the model. It should be noted that although environmental transmission now accounts for the majority of infections, there are more infections occurring from direct transmission on the first day. On subsequent days environmental contamination has built up and contributes more to transmission, with the remaining animals quickly becoming infected. The distribution of infection times is similar to that shown in Fig. 4 and thus not shown here. The median infection times for direct and environmental transmission are 2.0 and 2.4 when *A* = 1 compared to 2.3 and 3.0 when *A* = *N*.

**Figure 7:**
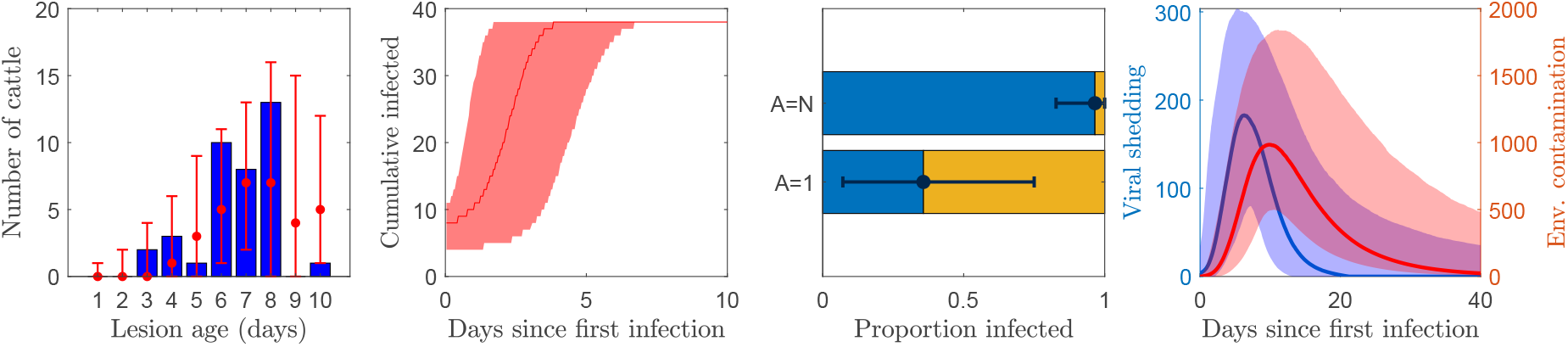
Dynamics of transmission for IP1b when the size of the environment is not considered (i.e. *A* = 1; cf. results for IP1b in Figs. 2, 3 and 5).

The simulated lesion ages when *A* = 1 are similar to both the observed ages (Fig. 7) and those for the model when *A* = *N* (cf. Fig. 2). Model selection as described in Section 2.3 between the models with *A* = 1 and *A* = *N* does not provide evidence in favour of one model over the other. Furthermore, when model selection is used to determine the best of the three models (combined transmission, direct only and environmental only) but with *A* = 1, there is no evidence to choose one model over another. This suggests that it is possible for environmental transmission alone to be the route of transmission if contamination rates are not affected by the size of the area being contaminated.

The median herd generation times (90% credible intervals) for viral shedding and environmental contamination are 7.78 (4.90-16.2) and 13.9 (10.9-20.3) respectively. These are almost identical to the previous results (Table 2). Similarly, the proportion of transmission before detection are similar to those shown in Fig. 6 (differences between medians were less than 0.02 for shedding and 0.01 for environmental contamination).

#### 3.5.2 Sensitivity to prior assumptions

To further assess model sensitivity, we fit the model to IP1b using uniform prior distributions for different combinations of parameters relating to transmission and environmental contamination. Here, we revert to using *A* = *N*.

First, we used uniform prior distributions for the two transmission parameters, *β_d_* and *β_e_*, both separately and together. The limits for the uniform distributions were chosen as the minimum and maximum of the samples used to the generate the informative prior distributions.

Posterior distributions for *β_d_* and *β_e_* are shown in Fig. 8. When a uniform prior distribution is used for a parameter, the median of the posterior distribution is larger than when using the informative prior. This is the case when either parameter has a uniform prior alone or both do together. Note that parameters not shown here did not change substantially from when informative priors were used (see Fig. S1).

**Figure 8:**
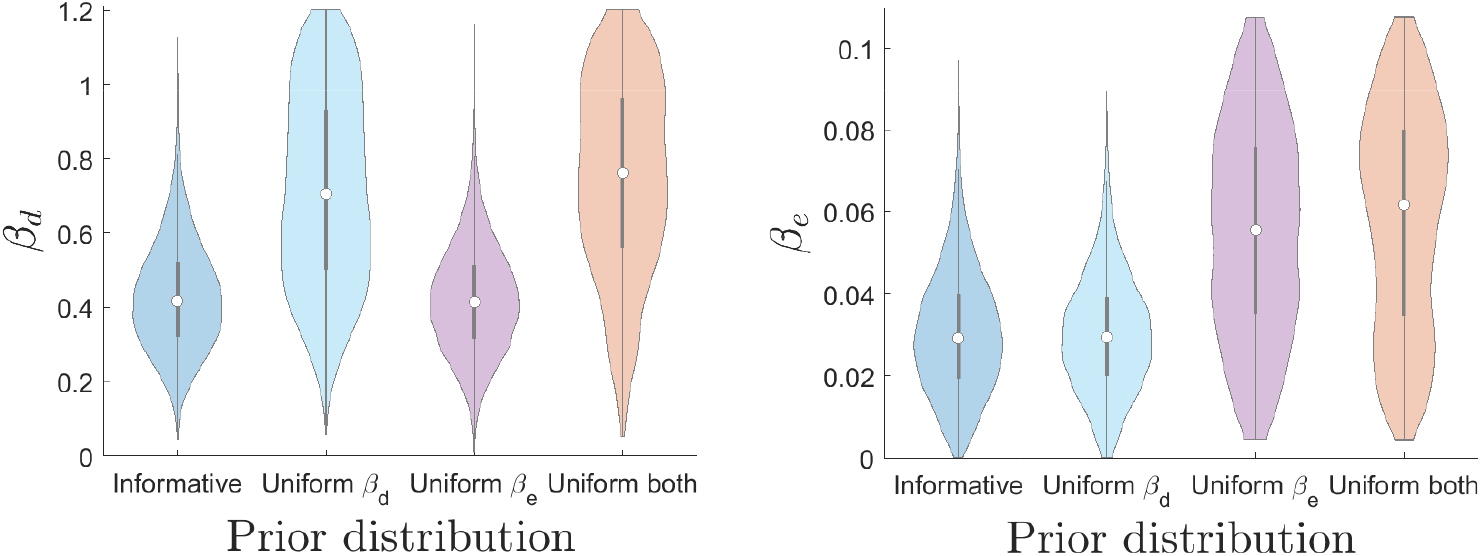
Posterior distributions for transmission parameters *β_d_* and *β_e_* for IP1b when fitted using different combinations of uniform prior distributions.

Second, we also considered uniform prior distributions for the contamination rates for the four areas of the environment, *α_i_, i* = 1, …4. In this case, the uniform distribution was bounded by 0 and the median of the informative prior distribution parameter multiplied by *A*. This gives the opportunity for contamination rates to be similar to when *A* = 1 (an increase of *A* cancels out the division by *A* in Eqn. 3) or similar to the model with *A* = *N* if the posterior distribution becomes close to that obtained using an informative prior distribution.

The posterior distributions for the contamination rates divided by the posterior median obtained using informative prior distributions are shown in Fig. 9. This allows us to see how the contamination rates relate to the environment size. The posterior distributions for each contamination rate remain fairly uniform but with small peaks that vary between 20 and 36 times higher than their respective informative prior medians. Other parameters not shown here did not change significantly from when informative priors were used (see Fig. S1).

**Figure 9:**
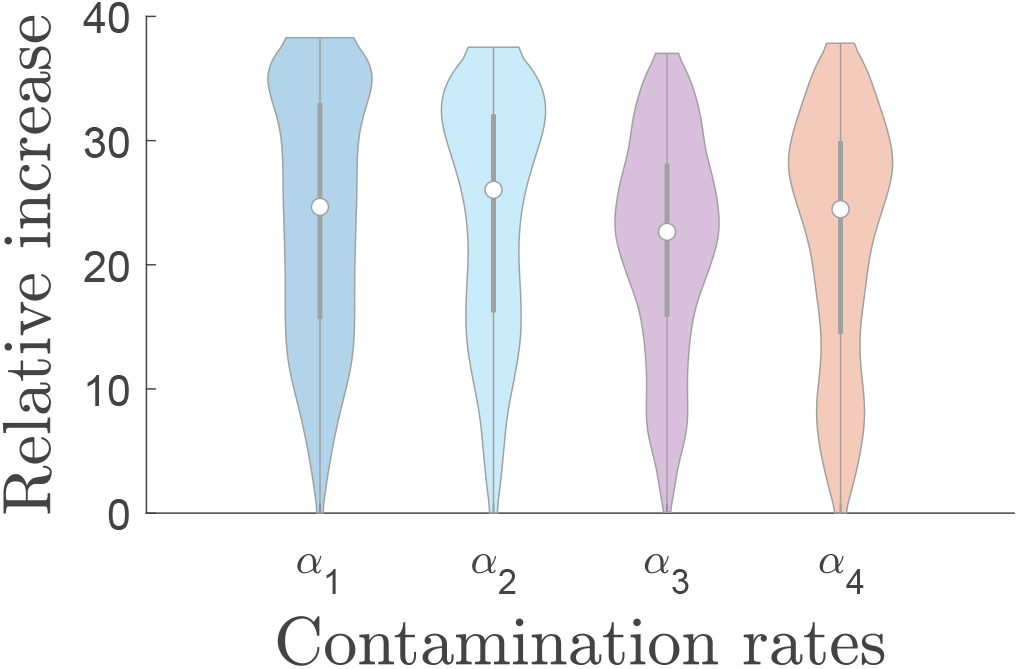
Posterior distribution for environmental contamination rates (*α_i_*) when using uniform priors relative to the posterior median when using informative priors. An increase by a factor of *N* (=38 for IP1b) is the equivalent of a contamination rate the same as when *A* = 1.

In all cases where non-informative priors were used, the summary transmission measures (*T_g_* and *θ*) were not substantially different to the original model. The median herd generation times have less than a 2.5% difference to the results shown in Table 2 and the proportion of transmission occurring before different detection times or thresholds were all within 0.025 of the results shown in Fig. 6.

## 4 Discussion

We have developed a model that shows the infectiousness profile of a farm for two different routes of transmission of FMDV. Our results show that although the contribution to transmission from environmental contamination is small, assuming contamination rates are affected by the size of the environment, it is the cause of infection of at least one animal in the majority of simulated outbreaks. Moreover, without control it can be a route of between-farm transmission and virus can remain in the environment after all animals have ceased to be infectious.

Parameters were estimated using ABC-SMC with prior distributions obtained using data from transmission experiments [10, 13]. There was some variation in the prior distributions for the different IPs, particularly in parameters closely relating to direct transmission and appearance of clinical signs. An increase in transmissibility, by increasing transmission or contamination parameters, may fit the data better. However, the changes when using informative priors were mostly small, indicating that the results from the transmission experiments are still appropriate for modelling an outbreak of FMD, at least for genetically similar strains of the virus.

The estimated initial number of infected animals for each IP, shown in Fig. 1, corresponds to other IPs during the 2007 epidemic which were thought to have had up to 15 initial infections [42]. The results are higher than might be expected based on the results found from the 2001 epidemic [1] (means of 4.29, 4.34, 4.33, 4.36 and 4.18 cattle infected initially compared to posterior means of 8.39, 6.15, 9.01, 4.40 and 2.12 respectively). However, some IPs were found to have many more initially infected animals during the 2001 epidemic [1] and models for 2001 may not be appropriate for the situation in 2007 where the epidemic was on a much smaller scale.

Infection proceeds quickly, with the whole herd typically infected within eight days (see Fig. 2). The simulated clinical signs broadly fit the pattern of the lesion age data, although it can overestimate the number of older lesions (IP1b and IP2a) and underestimate the days with a high number of new lesions (IP2a and IP3b). This can be partially explained by uncertainty in the accuracy of lesion ages. The margin of error in lesion age estimation is usually plus or minus one day but increases with lesions older than five days [42]. We did not incorporate uncertainty of lesion ages into the model but would not expect this to have a substantial effect on the inferences.

The model results clearly show that direct contact drives transmission; approximately 80% of all animals being infected via the direct route (Fig. 3). One reason for this is that the direct transmission parameter is larger than the environmental transmission parameter by a factor of 10 (see Table S2). Another contributing factor is that peak environmental contamination occurs later than peak viral shedding and, as can be seen in Fig. 2 and Fig. 4, the majority of animals become infected within the first 6 days. This leaves fewer susceptible animals at the time of peak environmental contamination, which occurs at approximately day 9. However, the environment remains infectious and still poses a risk of transmission between farms.

Model selection using ABC-SMC did not provide evidence to prefer either the combined or the direct transmission only model. This is probably due to the comparatively small impact on transmission from environmental contamination, and therefore the indirect impact on the measure being fitted (the number of animals with clinical signs). This also explains why model selection provided strong evidence against the environmental transmission only model. It should also be noted that the model selection algorithm favours models with fewer parameters, as is the case with the direct transmission only model in particular, as the higher the number of parameters, the smaller the probability that the perturbed particle is accepted [50].

Results show that the herd generation time is larger for environmental contamination than for viral shedding (see Table 2). We can compare this to generation time estimates from past epidemics to infer which route of transmission was causing infection between farms. Estimates of the mean herd generation time in the 2001 UK epidemic were 7.6 [52] and 6.3 days [25], which is slightly below our estimates based on viral shedding. However, the distribution of generation intervals (Fig. 1b in [25]), which has a large peak at 7 days, also has a small ‘bump’ at approximately 15 days. Although this could be incidental, it could be indicative of a separation between routes of infection, with a small proportion of later infections that occur via environmental contamination. Other estimates of the herd generation time for FMDV usually fall into the range of 5-15 days, with most instances being between 7-10 days [16, 28, 51]. Our median herd generation times based on viral shedding are within this interval, while those based on environmental contamination fall at the larger extreme of the interval.

The importance and the difference in the time scales of infection routes is also demonstrated by the proportion of transmission that occurs before certain times (see Fig 6). The times were chosen to demonstrate the impact of applying control measures early and were loosely based on other estimations of detection and control of FMD. The mean time from infection to detection in the 2001 UK epidemic was estimated to be approximately 8 days [12]. In our model, clinical signs first appear on average three days post introduction, so the corresponding detection time would be five days after clinical onset. Upon detection, some measures can be applied immediately, such as isolation, whereas the slaughter of animals will occur after a delay. The time from detection to slaughter can vary greatly between farms but was estimated to be on average close to one day in the later stages of the 2001 UK epidemic when response times were quickest [46, 52].

If we make this assumption, the very left on the top two plots in Fig 6 consider scenarios where detection would have occurred before any clinical signs. In the bottom two figures, we assume that detection occurs immediately after a certain percentage of the population have clinical signs and slaughter happens one day later. This is an approach used by previous modelling studies to approximate detection times [7, 9]. Nevertheless, this is a simplification of real infection to detection and detection to slaughter times. In reality there will be heterogeneity between individual farms and some may experience longer delays, changing the shape of the outbreak [46]. Therefore it is useful to consider different scenarios to show the range of results.

When considering between farm transmission via fomites, the level of environmental contamination is the main concern. Applying control measures at any time will substantially reduce the infectiousness from environmental contamination of an IP. Approximately 80% of the potential infectiousness will have been prevented if control is applied five days after the first clinical signs. For viral shedding, where infection may occur when moving animals or long distance via the wind, more urgent action is required. In this case, 80% of infectiousness is prevented if action is taken two days after the first appearance of clinical signs, though after five days less than half of infectiousness is prevented. There is some slight variation in *θ_S_* and *θ_E_* between IPs, possibly due to the herd size and the number of initially infected animals. A larger herd with few initial infected animals will have a longer outbreak than a small herd with many initial infections and therefore a higher proportion of infectiousness will occur after animals begin to show clinical signs.

To scale from transmission experiments to the field, we have assumed that the rate of environmental contamination is inversely proportional to the size of the environment, which we assumed to be linearly proportional to the herd size. If the environmental size is ignored, the model can still fit the 2007 epidemic data. In this case, the levels of viral contamination are approximate 20 times higher and environmental transmission contributes more than direct transmission. However, this does not seem to have an impact on the summary measures, namely the herd generation times and the proportion of infectiousness at different times post infection.

It is clear from the sensitivity analysis of the environment size and prior assumptions that an increase in transmission through increasing transmission parameters or contamination rates is preferred by ABC-SMC when fitting to the 2007 UK epidemic data. This is a consequence of the rapid spread of FMDV, such that an entire herd can be infected in a short period of time. Consequently, it is difficult to estimate an upper bound for parameters that increase transmission without prior information. This explains the broad distributions in Figs. 8 and 9 when using uniform prior distributions.

This study has used ABC-SMC to assess routes of transmission. We have shown that the parameters inferred from transmission experiments reflect transmission in the field, at least for a similar strain. We have also demonstrated the differences between direct transmission and transmission via environmental contamination of FMDV within a farm. Environmental transmission may play a small role within a farm, but virus remains in the environment after all animals have stopped being infectious and, hence, poses a danger of fomite transmission between farms and also a risk of re-suspension of virus into the air. Environmental contamination also presents an opportunity for another mode of detecting infected farms. Detection via clinical inspection requires veterinarians to attend a suspected infected site and the time taken to confirm presence of FMDV can be variable. Environmental sampling, as proposed by [6, 14, 15], may therefore provide an efficient alternative for sites that are at risk.

## Acknowledgements

This work was funded by the Department for Environment, Food and Rural Affairs (Defra) (grant code: SE2722). DS and SG acknowledge additional funding from the Biotechnology and Biological Sciences Research Council (BBSRC) (grant codes: BB/E/I/00001717, BB/E/I/00007036 and BB/E/I/00007037).

