## Supplementary Information for "Inferring transmission routes for foot-and-mouth disease virus within a cattle herd using approximate Bayesian computation"

Code for ABC-SMC and the transmission model is available online at
`https://github.com/DrJREllis/Inferring-transmission-routes-FMDV`

#### 1.1 ABC-SMC

1. Start at round t=1. Initialise the tolerance for the goodness-of-fit metric, $\epsilon_1$.
2. Generate a particle (i.e. set of parameters), $\theta_{i,t}$:
   (a) if $t = 1$, sample $\theta^{*}*$ independently from the joint prior density, $\pi(\theta)$;
   (b) if $t > 1$, sample $\theta^*$ from the previous population $\theta_{t-1}^{(i)}$ with weights $w_{j,t-1}$ generated during the previous round; perturb the particle to obtain $\theta^{**} \sim K_t(\theta|\theta^*)$ where $K_t$ is the perturbation kernel.
3. Simulate a candidate dataset $x^* \sim f(x|\theta^{**})$
4. Compare the simulated and observed data using the goodness-of-fit metric, $d(x^*, x_0)$. If $d < \epsilon_t$, accept the particle; otherwise (i.e. if $d \geq \epsilon_t$), go back to step (2).
5. Set $\theta_t^{(i)} = \theta^{**}$ and calculate the weight for the particle,
   (a) if $t = 1$, $w_{i,t} = 1$;
   (b) if $t > 1$, the weight is given by,
   $$w_{i,t} = \frac{\pi(\theta_{i,t})}{\sum_{j=1}^{N} w_{j,t-1} K(\theta_{i,t}|\theta_{j,t-1})}, \tag{1}$$
   where $\pi(\theta_{i,t})$ is the prior probability of the particle and $w_{j,t-1}$ is the weight of particle $j$ in the previous round.
6. Repeat steps (2)-(5) until 10,000 particles have been accepted.
7. Normalise the particle weights, so that they sum to one.
8. Calculate a new tolerance $\epsilon_{t+1}$ from the median value of the distances, $d$, for the round.
9. Increase the round number to $t + 1$ and go to step (2).

#### 1.2 ABC-SMC for model selection

When using ABC-SMC for model selection, the model $m$ is sampled first and then a particle $\theta$ is subsequently sampled for the chosen model.

1. Start at round t=1. Initialise the tolerance for the goodness-of-fit metric, $\epsilon_1$.
2. Generate a particle (i.e. set of parameters), $\theta_{i,t}$:
   (a) if $t = 1$, sample $(m^{**}, \theta^{**})$ from the prior distribution, $P(m, \theta)$;
   (b) if $t > 1$, sample $m*$ with probability $P_{t-1}(m^*)$ and perturb to obtain $m^{**} \sim KM_t(m|m^*)$ where $KM_t$ is the model perturbation kernel.
   Sample $\theta^*$ from the previous population $\theta(m^{**})_{t-1}$ with weights $w_{t-1}$ and perturb to obtain $\theta^{**} \sim KP_{t,m^{**}}(\theta|\theta^*)$ where $KP_{t,m^{**}}$ is the particle perturbation kernel for model $m^{**}$.
3. Simulate a candidate dataset $x^* \sim f(x|m^{**}, \theta^{**})$
4. Compare the simulated and observed data using the goodness-of-fit metric, $d(x^*, x_0)$. If $d < \epsilon_t$, accept the particle; otherwise (i.e. if $d \geq \epsilon_t$), go back to step (2).
5. Set $(m_t^{(i)}, \theta_t^{(i)}) = (m^{**}, \theta^{**})$ and calculate the weight for the particle,
   (a) if $t = 1$, $w_{i,t} = 1$;
   (b) if $t > 1$,
   $$w_{i,t} = \frac{P(m_t^{(i)}, \theta_t^{(i)})}{S}, \tag{2}$$
   Where
   $$S = \left( \sum_{j=1}^{\mathcal{M}} P_{t-1}(m_{t-1}^{(j)}) KM_t(m_t^{(i)}|m_{t-1}^{(j)}) \right) \left( \sum_{k;m_{t-1}=m_t^{(i)}} \frac{w_{t-1}^{(k)} KP_{t,m_t^{(i)}}(\theta_t^{(i)}|\theta_{t-1}^{(k)})}{P_{t-1}(m_{t-1} = m_t^{(i)}} \right), \tag{3}$$
   and $\mathcal{M}$ is the number of candidate models.
6. Repeat steps (2)-(5) until 10,000 particles have been accepted.
7. Normalise the particle weights and take the sum to obtain marginal model probabilities,
   $$P_t(m_t = m) = \sum_{i;m_t^{(i)}=m} w_t^{(i)}(m_t^{(i)}, \theta_t^{(i)}). \tag{4}$$
8. Calculate a new tolerance $\epsilon_{t+1}$ from the median value of the distances, $d$, for the round.
9. Increase the round number to $t + 1$ and go to step (2).

### 2 Inferring routes of transmission of foot-and-mouth disease virus within a cattle herd: Supplementary figures and tables

Table S1: Estimated age of lesions from clinical investigations during the 2007 UK FMD outbreak, data copied from (Ryan et al., 2008).

| IP | Herd size | Total infected | Age of lesions (days) 1 | 2 | 3 | 4 | 5 | 6 | 7 | 8 | 9 | 10 |
|---|---|---|---|---|---|---|---|---|---|---|---|---|
| IP1b | 38 | 38 | 0 | 0 | 2 | 3 | 1 | 10 | 8 | 13 | 0 | 1 |
| IP2a | 49 | 44 | 2 | 13 | 8 | 5 | 7 | 8 | 1 | 0 | 0 | 0 |
| IP3b | 47 | 36 | 3 | 17 | 11 | 2 | 3 | 0 | 0 | 0 | 0 | 0 |
| IP4b | 54 | 54 | 0 | 0 | 0 | 0 | 2 | 19 | 13 | 12 | 3 | 5 |
| IP7 | 16 | 14 | 1 | 3 | 4 | 4 | 2 | 0 | 0 | 0 | 0 | 0 |

Table S2: Parameter values (median and 90% credible intervals) used in the model and their sources.

| | Parameter | Prior | IP1b | IP2a | IP3b | IP4b | IP7 |
|---|---|---|---|---|---|---|---|
| $s_V$ | Shape parameter of log peak titre | 15.8 (6.88-35.6) | 19.1 (3.48-42.7) | 19.5 (3.79-43.7) | 16.7 (2.43-40.2) | 17.5 (2.71-41.4) | 17.7 (2.71-42.7) |
| $\mu_V$ | Mean of log peak titre | 9.67 (8.53-11.1) | 9.97 (8.31-11.7) | 9.79 (8.13-11.5) | 10.1 (8.36-11.7) | 10.3 (8.69-11.9) | 9.97 (8.23-11.8) |
| $\mu_{T_p}$ | Mean of log time of peak titre | 1.41 (1.22-1.61) | 1.34 (1.09-1.59) | 1.38 (1.12-1.62) | 1.34 (1.1-1.61) | 1.3 (1.05-1.57) | 1.35 (1.07-1.6) |
| $\sigma_{T_p}$ | Standard dev. of log time of peak titre | 0.4 (0.27-0.592) | 0.398 (0.214-0.631) | 0.413 (0.22-0.632) | 0.423 (0.233-0.65) | 0.443 (0.235-0.672) | 0.413 (0.221-0.641) |
| $s_{\gamma_g}$ | Shape parameter of viral growth rate | 2.23 (1.14-4.45) | 2.43 (0.415-6.21) | 2.53 (0.462-6.28) | 2 (0.212-5.72) | 2.49 (0.536-6.06) | 1.96 (0.149-5.89) |
| $\mu_{\gamma_g}$ | Mean viral growth rate | 8.21 (5.78-12.3) | 7.45 (3.2-12.8) | 7.59 (3.23-12.9) | 6.8 (2.34-12.4) | 6.91 (3.13-11.9) | 7.09 (2.2-12.6) |
| $s_{\gamma_d}$ | Shape parameter of viral decay rate | 5.29 (1.64-19.7) | 7.96 (0.958-22.7) | 7.62 (0.838-23.9) | 7.46 (0.906-21.8) | 7.77 (0.929-22) | 8.24 (1.07-26.6) |
| $\mu_{\gamma_d}$ | Mean viral decay rate | 2.74 (2.15-3.7) | 2.72 (1.14-4.37) | 2.71 (1.11-4.41) | 2.7 (1.12-4.38) | 2.76 (1.12-4.59) | 2.78 (1.18-4.51) |
| $\mu_{T_c}$ | Mean of log time of clinical onset | 1.32 (1.17-1.47) | 1.31 (1.1-1.54) | 1.32 (1.1-1.54) | 1.37 (1.16-1.57) | 1.24 (1.02-1.48) | 1.34 (1.13-1.54) |
| $\sigma_{T_c}$ | Standard dev. of log time of clinical onset | 0.32 (0.233-0.46) | 0.319 (0.222-0.425) | 0.328 (0.208-0.474) | 0.394 (0.287-0.526) | 0.204 (0.104-0.319) | 0.336 (0.215-0.473) |
| $\rho_{pc}$ | Correlation coefficient for $T_p$ and $T_c$ | 0.623 (0.082-0.867) | 0.584 (-0.055-0.922) | 0.597 (0.036-0.93) | 0.551 (0.078-0.912) | 0.595 (0.095-0.927) | 0.544 (0.025-0.903) |
| $\beta_d$ | Direct transmission rate | 0.315 (0.159-0.559) | 0.416 (0.209-0.687) | 0.37 (0.17-0.638) | 0.415 (0.195-0.697) | 0.572 (0.345-0.833) | 0.433 (0.214-0.712) |
| $\beta_e$ | Environmental transmission rate | 0.0273 (0.013-0.052) | 0.0285 (0.0074-0.0563) | 0.0303 (0.008-0.0603) | 0.0288 (0.0068-0.0574) | 0.0307 (0.0088-0.0576) | 0.0309 (0.0082-0.0617) |
| $\alpha_1$ | Floor contamination rate | 0.15 (0.122-0.185) | 0.15 (0.112-0.19) | 0.152 (0.111-0.193) | 0.151 (0.11-0.191) | 0.151 (0.112-0.194) | 0.149 (0.11-0.191) |
| $\alpha_2$ | Wall contamination rate | 0.0972 (0.077-0.123) | 0.0984 (0.069-0.13) | 0.0971 (0.068-0.128) | 0.0944 (0.067-0.126) | 0.1 (0.07-0.128) | 0.0954 (0.066-0.129) |
| $\alpha_3$ | Trough contamination rate | 0.326 (0.226-0.484) | 0.335 (0.169-0.525) | 0.329 (0.165-0.529) | 0.322 (0.161-0.517) | 0.344 (0.176-0.519) | 0.331 (0.17-0.535) |
| $\alpha_4$ | Faeces contamination rate | 0.373 (0.301-0.463) | 0.368 (0.264-0.48) | 0.374 (0.27-0.481) | 0.355 (0.255-0.475) | 0.373 (0.264-0.487) | 0.369 (0.271-0.488) |
| $\delta_1$ | Floor viral decay rate | 0.121 (0.088-0.152) | 0.121 (0.081-0.162) | 0.12 (0.082-0.16) | 0.122 (0.081-0.159) | 0.121 (0.081-0.163) | 0.12 (0.081-0.158) |
| $\delta_2$ | Wall viral decay rate | 0.131 (0.093-0.169) | 0.131 (0.084-0.175) | 0.132 (0.083-0.178) | 0.127 (0.081-0.173) | 0.133 (0.086-0.177) | 0.127 (0.077-0.185) |
| $\delta_3$ | Trough viral decay rate | 0.239 (0.175-0.304) | 0.238 (0.16-0.32) | 0.238 (0.163-0.311) | 0.246 (0.162-0.323) | 0.247 (0.166-0.323) | 0.24 (0.166-0.322) |
| $\delta_4$ | Faeces viral decay rate | 0.174 (0.14-0.209) | 0.175 (0.13-0.215) | 0.172 (0.131-0.217) | 0.172 (0.133-0.215) | 0.172 (0.121-0.215) | 0.174 (0.133-0.219) |
| $I_0$ | Initial number infected | U(1,N) | 8 (5-11) | 6 (4-9) | 9 (3-12) | 4 (2-6) | 2 (1-4) |

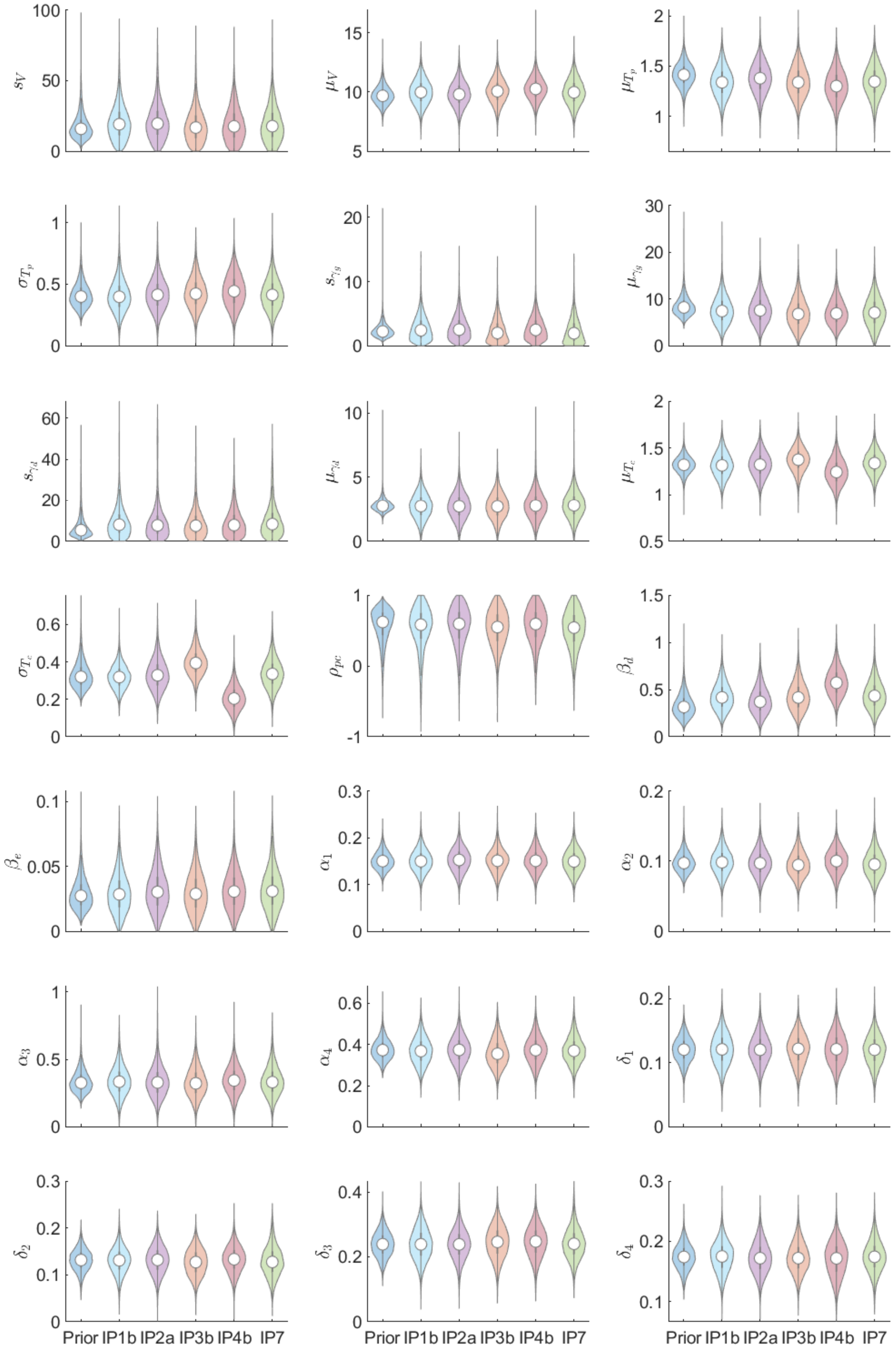


Figure S1: Prior and posterior distributions for within-host and transmission parameters for five farms infected with foot-and-mouth disease virus in the 2007 UK epidemic. Parameters are the shape parameter and mean of log peak titre, $s_V$ and $\mu_V$, viral growth rate, $s_{\gamma_g}$, $\mu_{\gamma_g}$, and viral decay rate $s_{\gamma_d}$, $\mu_{\gamma_d}$, the mean and standard deviation of log time to peak titre and clinical onset and the correlation coefficient $\mu_{T_p}$, $\sigma_{T_p}$, $\mu_{T_c}$, $\sigma_{T_c}$, $\rho_{pc}$, the transmission parameters $\beta_d$, $\beta_e$, and the environmental contamination and decay rates, $\alpha_i$, $\delta_i$.

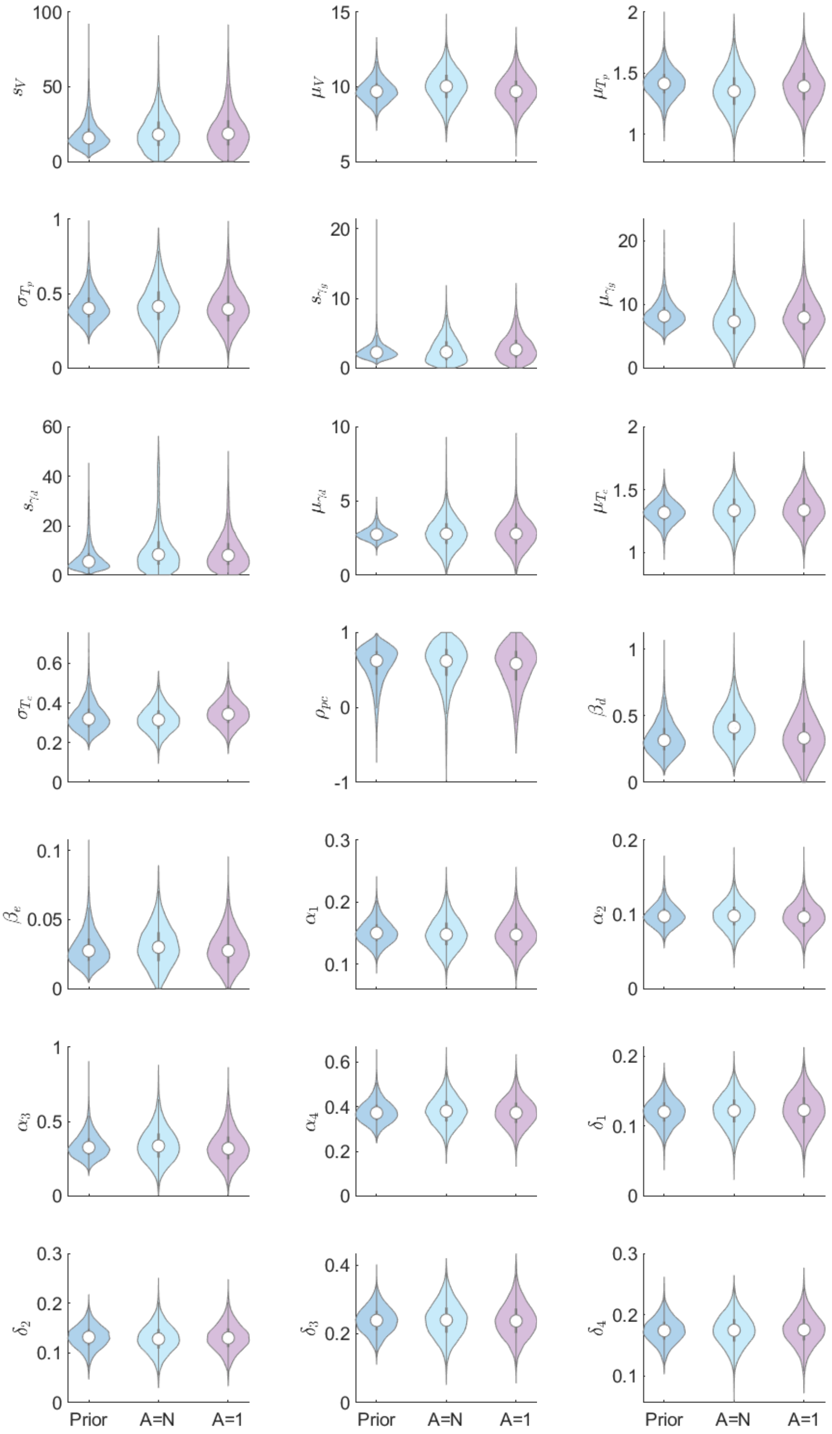


Figure S2: Prior and posterior distributions of the parameters from IP1b when the size of the environment is $(A = N)$ and is not $(A = 1)$ considered in the model.
